# Effect of Saudi and Egyptian pomegranate polyphenols in regulating the activity of PON1, PON2 and lipid profile for preventing coronary heart disease

**DOI:** 10.1101/570838

**Authors:** Mona Nasser BinMowyna, Manal Abdulaziz Binobead, Nawal Abdullah Al Badr, Sahar Abdulaziz AlSedairy, Islam Abdul Rahim Elredh, Wedad Saeed Al-Qahtani

## Abstract

Saudi and Egyptian pomegranate juice (APJ and EPJ) contains potent polyphenols antioxidants which were found to reduce serum and macrophage oxidative stress. The protective effects of APJ and EPJ against atherogenicity were evaluated by feeding mice with hypercholesterolemic diet supplemented with PJ. PJ consumption reduced mice serum Triglycerides (TG), Total cholesterol (TC) and (low density lipoprotein (LDL-c) and increase in the High-density lipoprotein (HDL-c) compared with mouse with control diet or atherogenic diet. The risk ratio and atherogenic index were significantly lower in either APJ or EPJ supplemented group. Paraoxonase 1 (PON1) which remains associated with HDL showed significant increase in the activity in mice supplemented with PJ compared to mice with atherogenic diet (AD). PJ consumption significantly reduced lipid peroxidation and increased glutathione levels. The reduction in lipid peroxidation varied from 57 to 76%. PJ supplementation reduced peritoneal macrophages (MPM) mediated oxidation of LDL by 68 to 82% and decreased mouse peritoneal macrophages (MPM) Ox-LDL uptake by 31 to 48%. A significant up regulation of macrophage PON2 activity was induced by PJ reducing the macrophage oxidative stress. In conclusion, PJ consumption resulted in anti-oxidative and antiatherogenic effects on serum and macrophages which could prevent atherosclerosis and coronary heart diseases.

## 1. Introduction

Over the last few years, the role of polyphenols in reducing risk of chronic diseases has received much attention. Polyphenols are naturally occurring phytochemicals contained in most of the fruits, vegetables, cereals and beverages (Grosso, 2018). Recent evidences suggested potential beneficial effects of polyphenols in regulating vascular and endothelial function by lowering blood pressure, improving endothelial function, increasing antioxidant and anti-inflammatory activities, inhibiting low-density lipoprotein oxidation, and reducing inflammatory responses (Giglio et al., 2018). They are associated with prevention and treatment of cardiovascular diseases (CVDs) reducing CVD-related mortality. One of the most evident biomarkers for CVD risk is high LDL (Low-density lipoprotein) cholesterol and low HDL (High-density lipoprotein) cholesterol (Hollands et al., 2018). Atherosclerosis the major cause of CVD is mediated by endothelial dysfunction caused due to accumulation of LDL. Endothelial dysfunction leads to infiltration of LDL and their subsequent oxidation to oxidized-LDL (ox-LDL) by interacting with superoxide anions (Moss et al., 2018). High oxidative stress in conditions such as CVD, reduces the serum concentration and activity of paraoxonase-1 (PON1) (Dizaji et al., 2018). Polyphenols reduces atherosclerosis by increasing HDL, reducing LDL and total cholesterol (Santhakumar et al., 2018).

Pomegranate (*Punica granatum* L.) has traditionally been used for treatment of many chronic diseases due to its multi-factorial beneficial effects. Pomegranate extracts are known to have anti-inflammatory, antidiabetic and antioxidant activity (AlMatar, 2018). Studies revealed that pomegranate is rich in polyphenols and includes anthocyanins and anthoxanthins. Ellagic acid and hydrolysable ellagitannins are the principal polyphenols component having antioxidant properties. Punicalagin, an ellagitannins found in pomegranate is known for its antiatherogenic activities (Les et al., 2018). It prevents accumulation of lipids in macrophages and foam cell formation (Atrahimovich et al., 2018). Other hydrolysable tannins include punicalain and gallic acid (Rosenblat et al., 2015).

Paraoxonase is an HDL-associated esterase that prevents oxidation of lipids in lipoproteins. Studies revealed that consumption of pomegranate juice (PJ) promotes binding of HDL to paraoxonase 1 (PON1), thereby increasing the hydrolytic and peroxidative activity of paraoxonase (Kaplan, 2001). PON1 is synthesized in the liver and transported in plasma in association with HDL. The decrease in the PON1 activity may lead to higher levels of cholesterol (Estrada-Luna et al., 2018). Other associated paraoxonase are PON2 and PON3. PON2 is expressed in most tissues including macrophages and plays a crucial role in preventing atherosclerosis. PON3 is found associated with HDL, mitochondria and endoplasmic reticulum (Moya and Manez, 2018). Studies suggested that pomegranate polyphenols reduce macrophage oxidative stress and formation of foam cells by over expression of macrophage PON2. PON2 prevent macrophage oxidation and oxidation of LDL by inhibiting the formation and release of reactive oxygen species (ROS) and reactive nitrogen species (RNS) (Aviram and Rosenblat, 2012).

The present study envisaged the effect of polyphenols of Saudi and Egyptian Pomegranate juice in regulating the activity of PON1, PON2, serum lipids and macrophage oxidative stress, and their role in preventing the coronary heart disease in mice.

## 2. Materials and methods

### Pomegranate fruits

Ripened fruits of the Saudi pomegranate were obtained from the local market of Taif and Egyptian pomegranate fruits from the local market of Riyadh city. The entire fruit was squeezed with a juicer and filtered through 15 μm membrane to remove insoluble residue. Fresh PJ was prepared daily during the entire experimental period.

### Experimental animals

Twenty-five CD1 mice were obtained from the experimental animal’s center, King Saud University. Mice weighing 25 ± 5 g were used in the study. The mice were housed in wire-bottomed cages and maintained in controlled environment (temperature of 22 ± 2 ° C, relative humidity of 50 ± 5%, and 12h light/dark cycle). Mice were divided into five groups, comprised of five mice per group. The first group was fed with normal control diet (CD), the second group with high-fat diet referred as atherogenic diet (AD), the third group was fed with high-fat diet supplemented with 200 μL Saudi pomegranate juice (APJ) containing 0.42 μmol of total polyphenols, the fourth groups was fed with high-fed diet supplemented with 200 μL Egyptian pomegranate juice (EPJ) containing 0.39 μmol of total polyphenols, the fifth group was fed with high-fat diet supplemented with 200 μL mixture of both APJ and EPJ. Supplementation was carried out for six month and diets was delivered directly to the stomach using oral gavage.

### Diets used in the experiment

The high-fat atherogenic diet contained (w/w) saturated fat 20%, cholesterol 1.5%, mineral salts mixture 2.5%, casein 20%, vitamin mixture 1%, cellulose 5% and starch 50%. The control group received water without PJ, saturated fat, cholesterol and casein. All the groups after supplementation had access to drinking water of 2-5 mL/day. Diets were stored at 4°C in plastic containers until used.

### Serum analyzes

At the end of the experiment, the mice were grafted and anesthetized using diethyl ether. The blood samples were then withdrawn and separated using a centrifuge. Serum analyses of total cholesterol, triglyceride, HDL-c, LDL-c were determined as previously described (Rosenblat et al., 2015).

### Serum lipid peroxidation

The extent of lipid peroxidation in plasma was determined by measurement of monodialdehyde (MDA) formation at 534 mm using the thiobarbituric acid reactive substances (TBARS) method. The reduced glutathione (GSH) in the plasma was estimated by its reaction with dithio-bis-2-nitrobenzoic acid (DTNB) that give a yellow coloured complex with maximum absorption at 412 nm.

### Serum PON1 activity

Serum PON1 activity was measured using phenyl acetate as the substrate (Gaidukov and Tawfik, 2005). Initial rates of substrate hydrolysis were determined spectrophotometrically at 270 nm. The assay mixture included 5 μL serum (diluted 1:3), 1.0 mmol/L phenylacetate, and 1 mmol/L CaCl_2_ in 50 mmol/L Tris HCL (pH 8.0). The results were calculated assuming the molar extinction coefficient of phenyl acetate to be 1,310 L mole^−1^ cm^−1^. One unit of arylesterase activity is equal to 1 μmoL of phenylacetate hydrolyzed/min/mL.

### Mouse peritoneal macrophages (MPM)

MPM were harvested from the peritoneal fluid of the mice, 3 days after IP injection into each mouse of 3mL of aged thioglycolate (20g/L) in saline. The cells (10-20 x10^6^ /mouse) were washed and centrifuged three times with phosphate-buffered saline (PBS) at 1000 x g for 10 min.

### MPM mediated oxidation of LDL

LDL was separated from plasma by density-gradient ultracentrigugation. MPN were incubated with LDL (100 μg of protein/mL) in the presence of 5 μM of CuSo_4_ for 5h at 37°C. LDL oxidation was measured by TBARS assay (Buege and Aust, 1978).

### MPM uptake of Ox-LDL

Ox-LDL was conjugated to fluoroisothiocyanate (FITC) for cellular lipoprotein uptake studies (Bass et al., 1983). MPM were incubated at 37°C for 3h with FITC-conjugated Ox-LDL at a final concentration of 25μg of protein/mL. The uptake of the lipoprotein was determined by flow cytometry. Cellular fluorescence was measured in terms of mean fluorescence intensity (MFI).

### MPM PON2 activity

Lactonase activity was measured in intact cells using dihydrocumarin (DHC) as a substrate. DHC (1mmoL/L) was added to the cells in 1mL of 1 mmol/L CaCl_2_ in 50 mmol/L of Tris HCL, pH 8.0. One unit of lactonase activity = 1 μmoL of DHC hydrolyzed/min. The absorbance was monitored at 270nm for 10 min after substrate addition.

### Statistical Analysis

Data obtained from the respective experiments were pooled and the means from these were used in statistical analysis using SPSS version-25. P values < 0.01 and < 0.05 were considered as significant. Means were separated with Duncan Multiple Range Test (DMRT) in respective tables and further verified using Akaike’s information criterion (AIC) and Hurvich and Tsai’s criterion.

## 3. Results and Discussions

The effect of hypercholesterolemic diets on different treated groups revealed that body weight gain significantly reduced in the group supplemented with APJ and EPJ compared to mice group with either control diet (CD) and atherogenic diet (AD). The reduction in net weight gain varies from 20.6 g to 22.4 g in APJ and EPJ supplemented diet (Fig.1). The result is in consistent with the earlier findings on reduction of body weight of mice treated with pomegranate seed oil and seed residue (Elbandy and Ashoush, 2012). However, more reduction in body weight was noticed in group treated with both APJ and EPJ supplemented diet (7.6 g). The result thus revealed that the dual effect causes more reduction in body weight compared to individual treatment with either APJ or EPJ alone. The least information criteria of net body weight gain according to Akaike’s information criterion (AIC) and Hurvich and Tsai’s criterion (AICC) was 812.851 (*R*^2^=0.42). Similarly, significant reductions in the relative heart weight were observed in the treated groups compared to control. The reduction in weight is 80.7% and 69.8% in APJ and EPJ supplemented mice group and 67.5% in APJ+EPJ supplemented mice group compared to AD group.

**Fig. 1.**
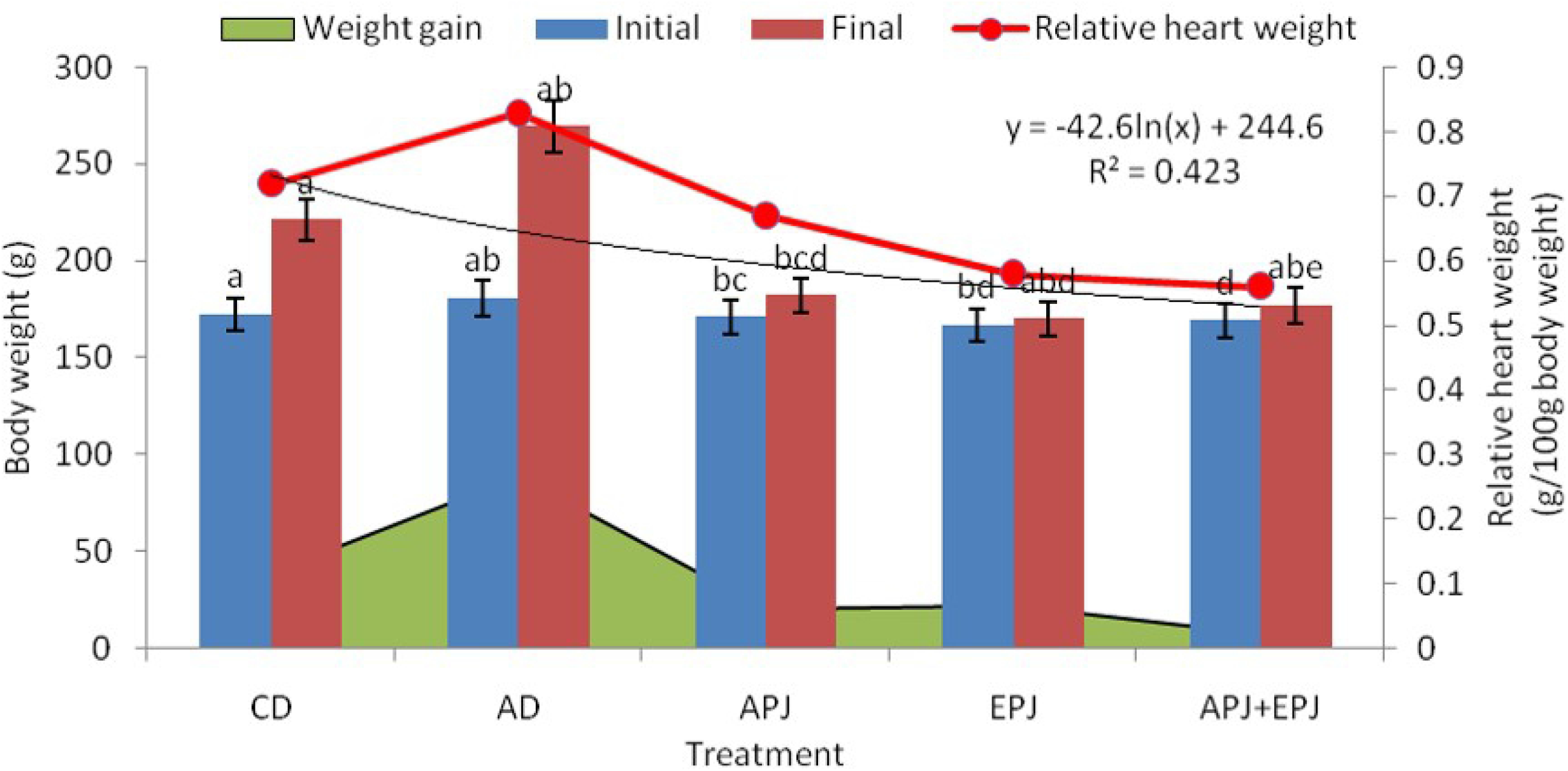
Variation in body weight and relative heart weight in high-.

The plasma lipid profile revealed significant higher level of TG (172.12 mg/dl), TC (153.34 mg/dl) and LDL-c (72.39 mg/dl) and lower level of HDL-c (33. 45mg.dl) in mice supplemented with atherogenic diet (AD). While mice supplemented with APJ, EPJ and combined APJ and EPJ diet showed significant lower level of TG, TC, and LDL-c and higher level of HDL-c compared to AD supplemented group (Fig.2). Thus, the present findings revealed decrease in the lipid fractions in PJ supplemented mice group, thereby reducing atherosclerotic process and chance of cardiovascular diseases (Berek and Bobinski, 2009). High HDL-c signifies anti-atherogenic properties and inhibition of LDL-oxidation. Similarly, the risk ratio (RR) and atherogenic index (AI) values the indicator of coronary and cardiovascular risk were significantly lower in mice supplemented with either APJ or EPJ or APJ and EPJ combined supplemented diets.

**Fig. 2.**
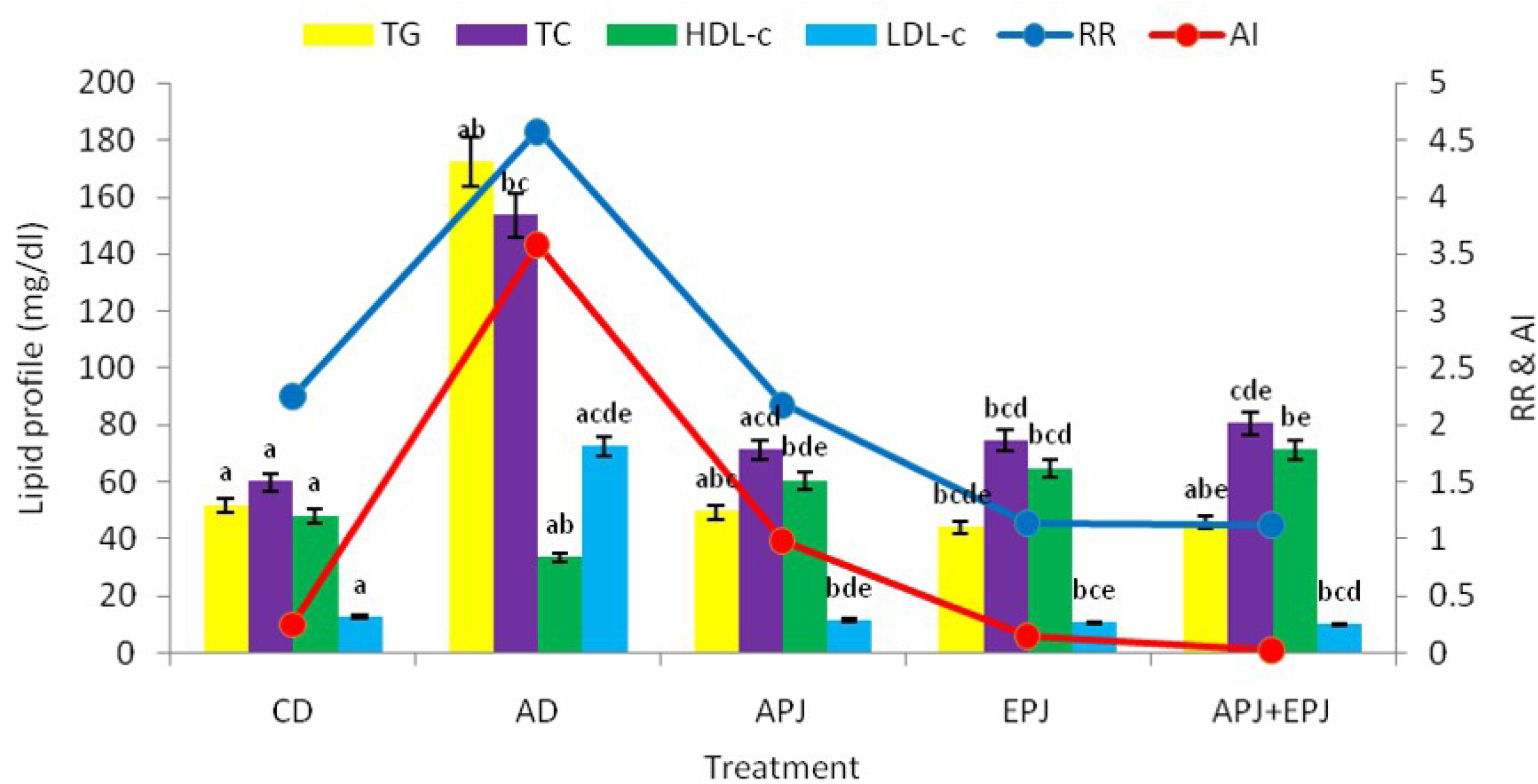
Lipid profile in high-fat diet fed mice supplemented with.

The present result revealed significant reduction in lipid peroxidation (MDA) in mice treated with either APJ or EPJ compared to mice treated with control diet (CD) and atherogenic diet (AD). The reduction varies from 57.1 to 76.3% in EPJ and APJ supplemented group (Fig.3). Thus, this is a clear indicative of effect of pomegranate juice in reduction of autocatalytic activities of lipid peroxidation thereby minimizing tissue damage. The reduction is more pronounced in group treated with both APJ and EPJ. Similarly, glutathione (GSH) and indicator of antioxidant activity, significantly increased in mice treated with APJ (47.2%) and EPJ (43.4%) as well as in combined application of APJ and EPJ (44.2%). In contrast, the level is significantly higher in mice with atherogenic diet. The reduction in MDA and enhancement of GSH level indicated that both APJ and EPJ have antioxidant properties capable of preventing tissue damage or cell death (*p*<0.05, *R*^2^=0.59).

**Fig. 3.**
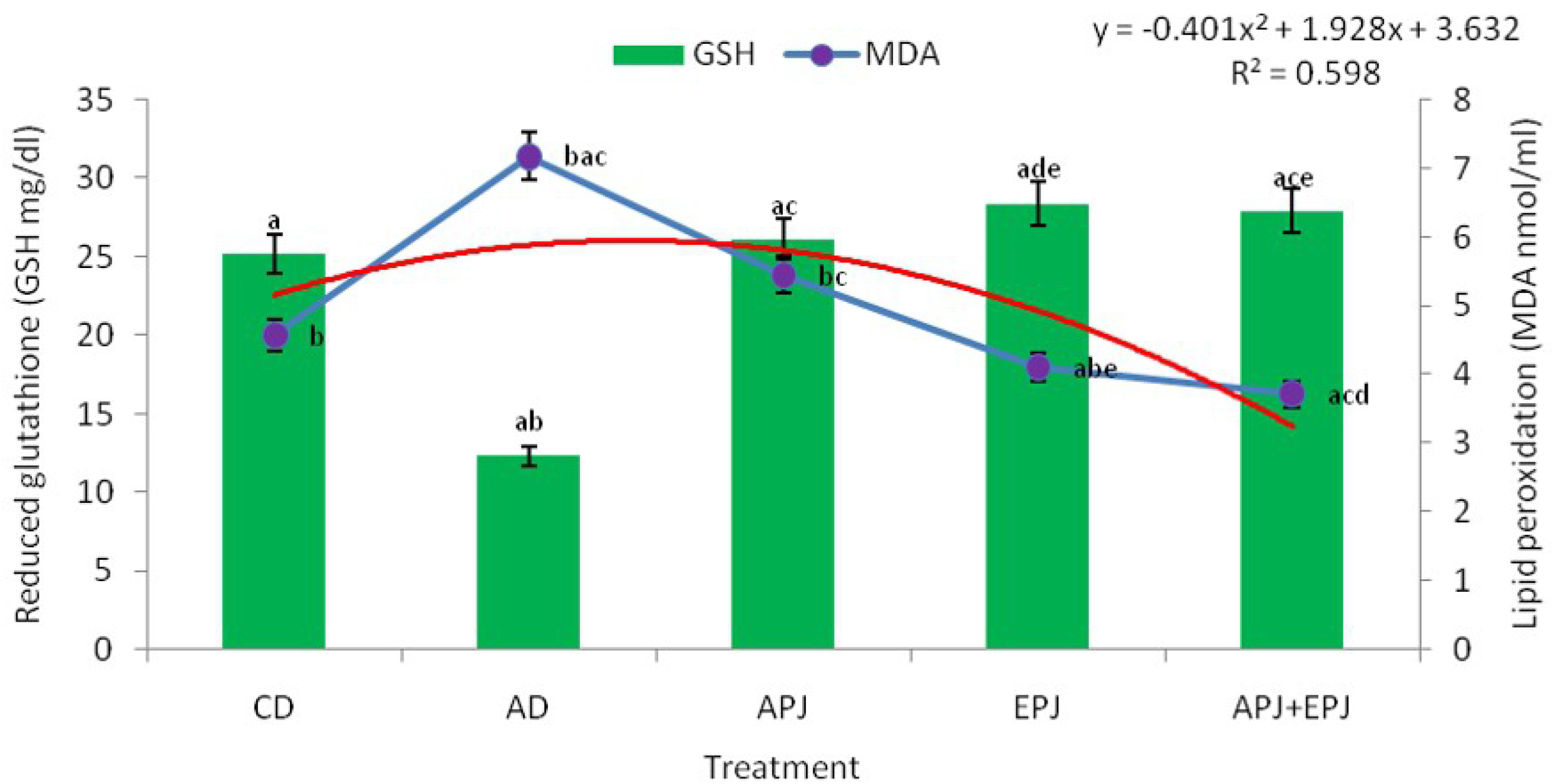
Plasma GSH and MDA in high-fat diet fed mice suppleme.

Serum paraoxonase (PON1) enzyme activity significantly decreased in mice supplemented with hypercholesterolemic diet compared to control and PJ treated (APJ, EPJ and APJ+EPJ) mice groups (Fig.4). The present result indicated that high-fat diet effects the paraoxonase activity increasing the risk of atherosclerosis resulting in coronary diseases (Estrada-Luna et al., 2018). There was marked increase in the paraoxonase activity in the mice group treated with PJ diet. The serum paraoxonase activity in PJ treated mice was 58% higher that mice with mice with atherogenic diet. This result is an indicative of high PON1 activity in PJ treated mice protecting LDL and HDL against oxidative processes, preventing the formation of atherogenic ox-LDL molecules (Zielaskowska and Olszewska, 2006). Research indicates that serum PON1 activity is inversely proportional to risk of atherosclerosis which confirms its protective properties in preventing cardiovascular diseases (Rosenblat et al., 2006).

**Fig. 4.**
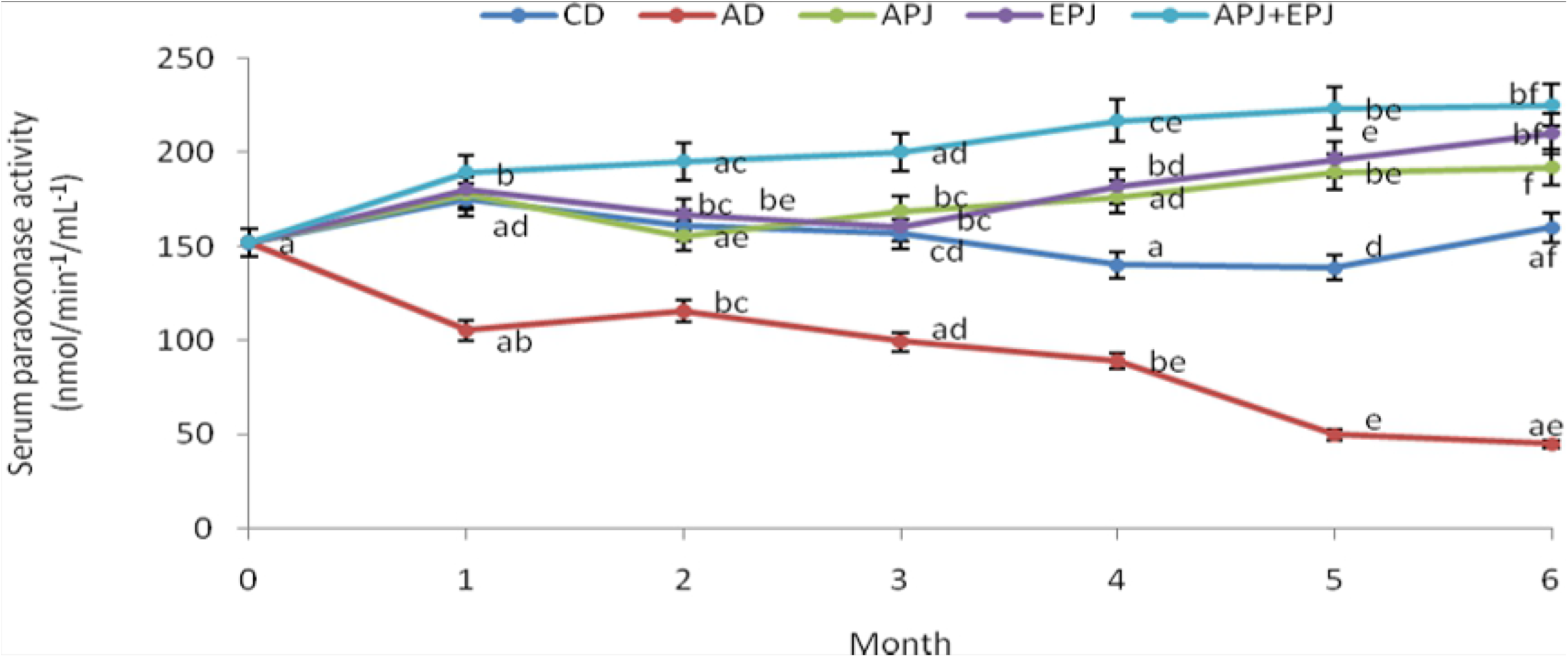
PON1 activity in high-fat diet fed mice supplemented with.

### MPM mediated oxidation of LDL

APJ, EPJ or APJ+EPJ supplemented mice groups showed significant decrement by 82.0 and 79.8%, 78.6 and 76.1%, 70.3 and 68.3% respectively in MPM mediated LDL oxidation compared to CD and AD supplemented mice groups (Fig.5). The result is in conformity with some of the earlier findings (Aviram and Rosenblat, 2004; Rosenblat et al., 2010). The result indicated that APJ was the most potent antioxidant against LDL oxidation. The decrement is significant with different PJ treatment as evidenced by correlation coefficient of *R*^2^=0.88, (*p*<0.05).

**Fig. 5.**
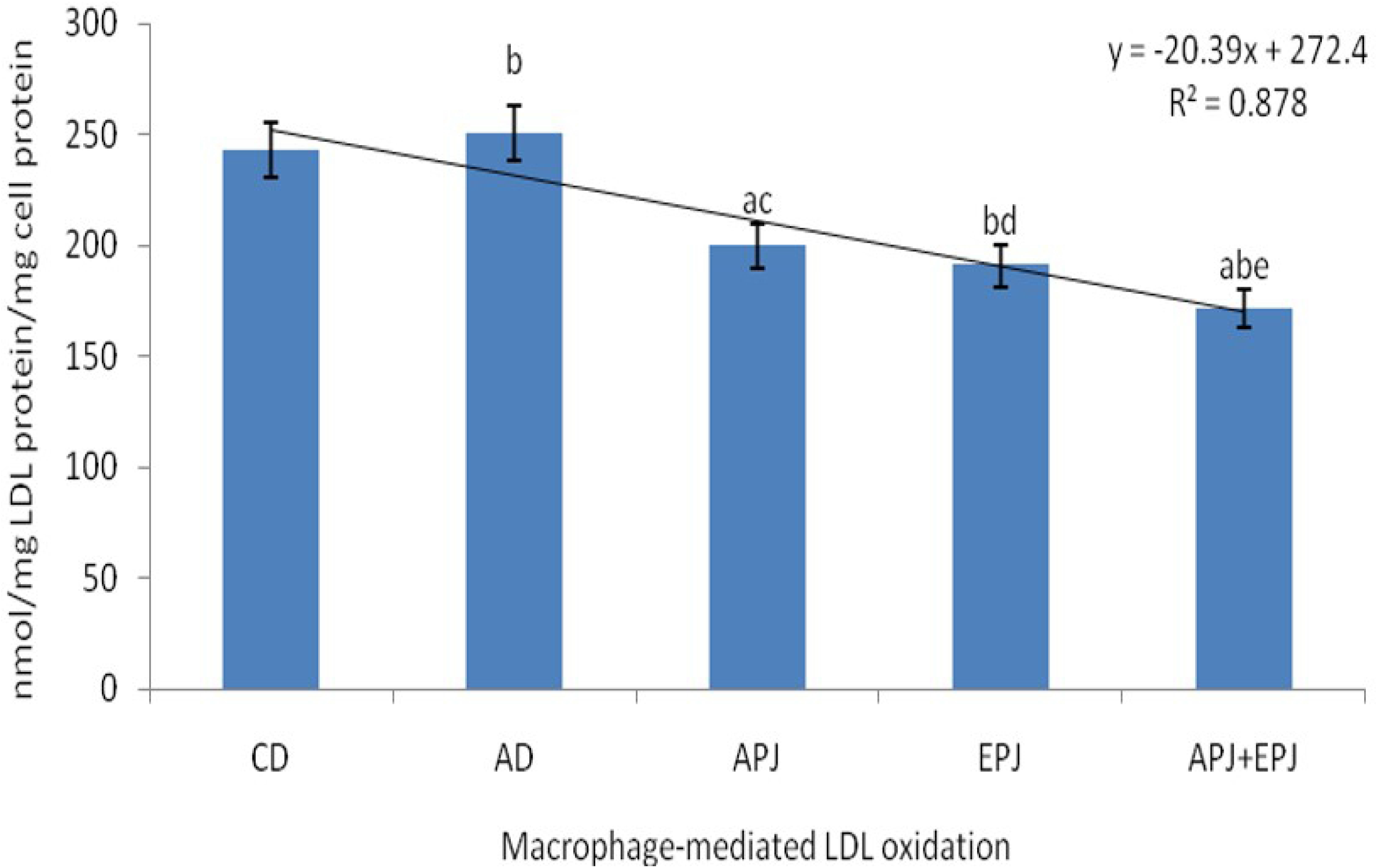
Macrophage mediated LDL oxidation in high-fat diet fe.

### MPM uptake of Ox-LDL

As evidenced from earlier studies oxidized LDL uptake by macrophages can lead to accumulation of macrophage cholesterol and formation of foam cell (Aviram and Rosenblat, 2004). In the present study, MPM from PJ treated mice group showed decreasing trend in Ox-LDL uptake compared to control and atherogenic group (Fig.6). The uptake was significantly decreased by 47.8% in MPM from APJ supplemented group, 46.5% from EPJ supplemented group and 36.7% from APJ + EPJ supplemented group. Similar result was obtained in some of the earlier studies (Rom et al., 2016).

**Fig. 6.**
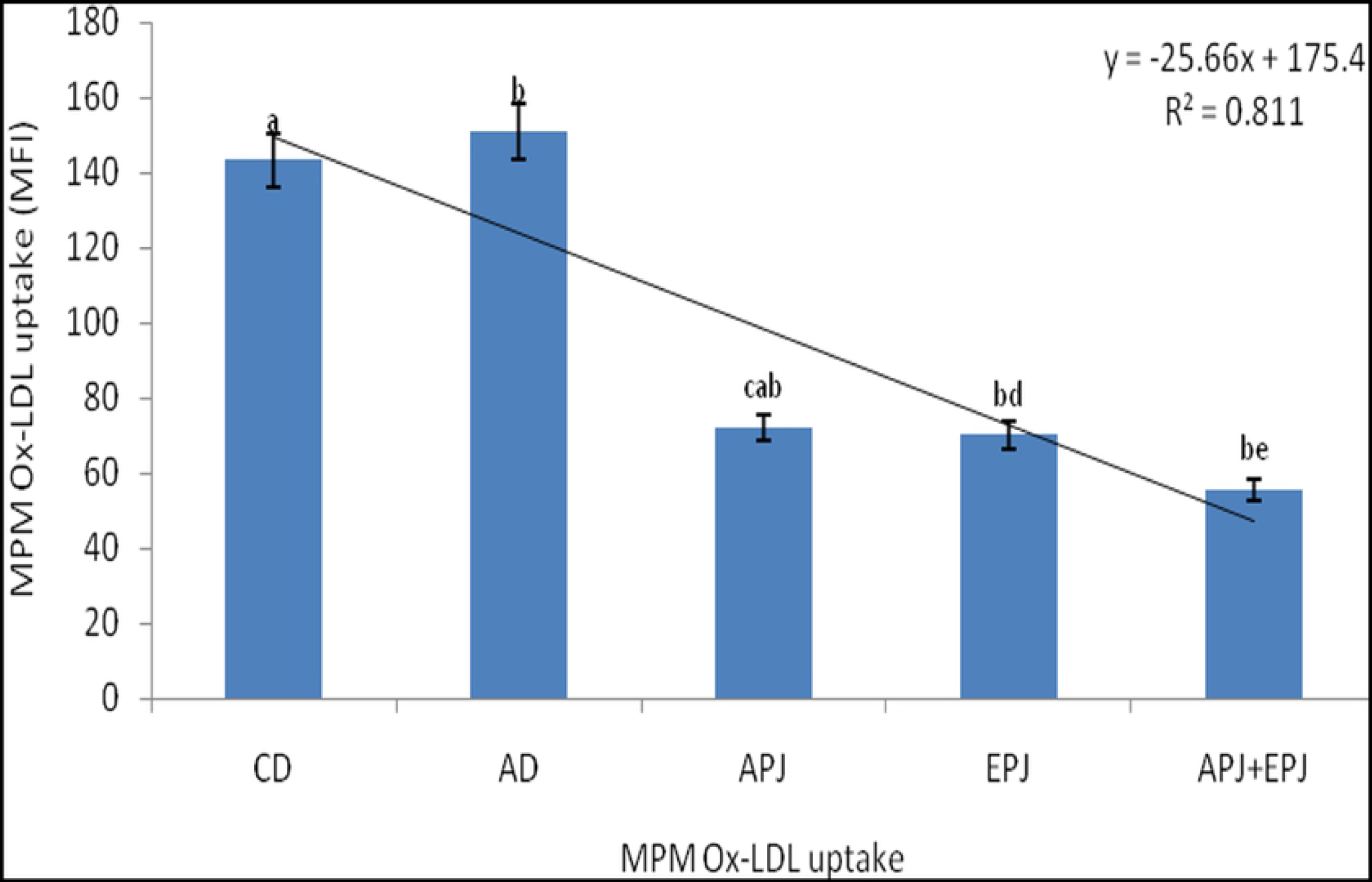
Macrophage mediated Ox-LDL uptake in high-fat diet fed.

MPM PON2 lactonase activity significantly increased by 54.6%, 50% and 48% in APJ, EPJ and APJ+EPJ supplemented mice group compared to group with atherogenic diet (Fig.7). The increase is significant with different PJ treatment as evidenced by correlation coefficient of *R*^2^=0.54, (*p*<0.05). The finding is pertinent to some of the earlier studies in which PJ was shown to increase in macrophage PON2 activity (Rosenblat et al., 2010). The increase is 2 to 3fold when compared to group with control diet. This increase may be attributed to increased PON2 relative gene expression following PJ supplementation. The present result indicates that PON2 may act as cellular antioxidant reducing oxidative stress through its anti-atherogenic role (Aviram and Rosenblat, 2004).

**Fig. 7.**
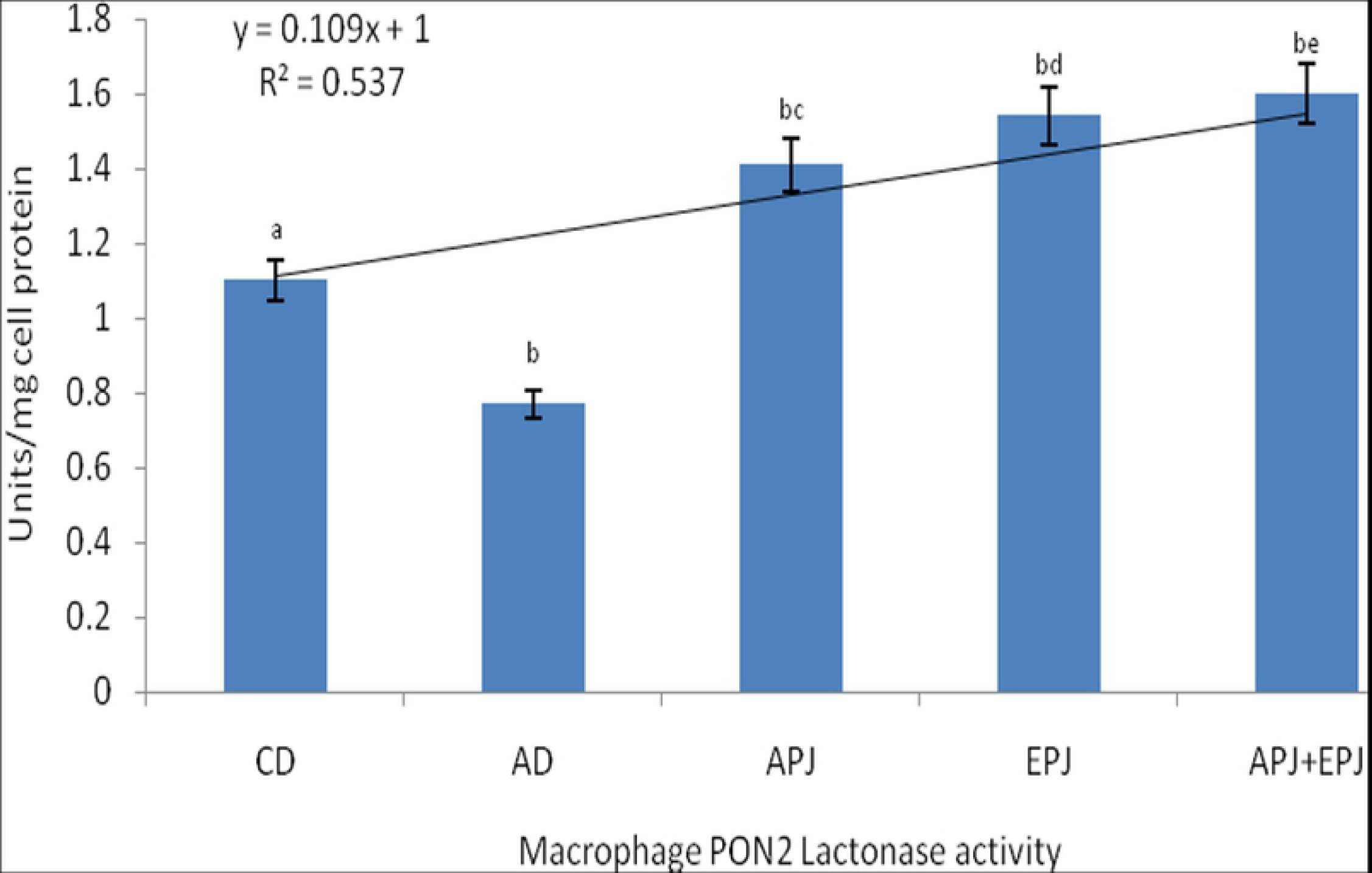
Macrophage PON2 activity in high-fat diet fed mice supple.

## 4. Conclusion

The present study reveals that dietry Saudi and Egyptian pomegranate juice significantly reduces oxidative stress by upregulating PON1 and PON2 activity and prevents LDL oxidation thus proofing its protective role in the progression of coronary heart diseases. The findings indicate that pomegranate juices can be preferred as a potential nutritional therapy in prevention of atherosclerosis and thereby overcoming the possible risk of developing heart diseases. This study highlighted the potential antioxidant effect of pomegranate polyphenols.

